# Incoherent feedforward motifs as immune change detectors

**DOI:** 10.1101/035600

**Authors:** Eduardo D. Sontag

## Abstract

We speculate that incoherent feedforward loops may be phenomenologically involved in self/nonself discrimination in immune-infection and immune-tumor interactions, acting as “change detectors”. In turn, this may result in logarithmic sensing (Weber phenomenon) and even scale invariance (fold-change detection).

## Introduction

A number of authors have proposed that lymphocytes mount a sustained response only when faced when a sufficiently fast increase in their level of stimulation (antigen presentation, proliferation rates of infected cells or tumors, stress signals). In contrast, even when a new motif triggers an immune response, its chronic presence may result in adaptation: downregulation or even complete termination of the inflammatory response. This dynamic feature is thought to complement discrimination mechanisms based on the self/nonself dichotomy.

One of the earliest such suggestions originated in work by Grossman and Paul [5], who in 1992 postulated the “tunable activation threshold” model for immune responses: effector cells in the innate or adaptive systems should become tolerant to continuously expressed motifs, or even gradually increasing ones, but should induce an effector response when a steep change is detected. Among notable recent variations upon this theme are the “discontinuity theory” postulated by Pradeu, Jaeger, and Vivier [10], and the “growth threshold conjecture” due to Arias, Herrero, Cuesta, Acosta, and Fernández-Arias [2]. (We will briefly discuss the mathematical models proposed in these references.) Evidence toward this role of dynamics include the examples of natural killer cells decreased activation under chronic receptor activation, endotoxin tolerance in macrophages, and B and T cell anergy. The tolerance of self -as well as non-self such as commensal bacteria– is consistent with this view, as is the tolerance of slow-growing tumors and even the treatment of allergies by slow desensitization through antigen exposure. This hypothesis remains to be tightened and experimentally verified. Obviously, the existence of autoimmune diseases argues against it, though Pradeu, Jaeger, and Vivier posit that possible explanations for chronic autoimmune disorders might be intermittent (as opposed to constant) exposure to antigens, or autoantigenic drift.

Our purpose in this note is not to argue pro or against the merits of the hypothesis, but merely to propose a very simple class of mathematical models that might achieve this type of dynamic discrimination. We provide a phenomenological “toy” model, and suggest possible biological mechanisms. Our model is based on incoherent feedforward loops (IFFLs), the incoherent path being implemented for example by natural Treg cells at a population level, or by ITIM/ITAM opposing controls at the intracellular level. We speculate that the change-detection features of IFFLs are involved in self vs non-self immune distinctions, thus complementing mechanisms such as kinetic proofreading. This phenomenology might help explain tolerance to self-antigens and immune reaction to both tumors and acute infections. More interestingly, our model predicts behaviors (Weber law, scale invariance) that could be in principle experimentally tested.

## Feedforward motifs

Several variants of the feedforward loop (FFL) “motif” (Fig. 1) describe instances of cell population interactions as well as of intracellular metabolic pathways, signaling networks, and genetic circuits [1]. FFLs play a role in systems ranging from bacterial chemotaxis and microRNA regulation to bacterial carbohydrate uptake via the carbohydrate phosphotransferase system, and also in mammalian cells as control mechanisms that regulate stress responses to free radicals, bacterial or viral infections, and cancer, and in the regulation of meiosis, mitosis, and post-mitotic functions in differentiated cells. They are statistically over-represented in natural systems (Fig. 1). In an incoherent feedforward loop (IFFL), an external cue or stimulus *x* activates or inactivates (represses) an intermediate molecular species *Y* which, in turn, activates or inactivates a downstream species *Z*. Through a different path, the signal *X* activates or inactivates the final species *Z*, in such a way that the net effect is opposite from that of the indirect path. This antagonistic (“incoherent”) effect endows the IFFL motif with powerful signal processing properties [1]. Specifically, an IFFL such as the “Type I” one shown in the bottom left panel of Fig. 1 has, for an appropriate mathematical model of interactions, a dynamical behavior that allows detection of change. Intuitively, a signal from the top node, *X*, will activate the bottom node, *Z*. An indirect path, through an auxiliary node *Y*, will -with some delay due to the signal processing by *Y*– eventually de-activate *Z*. Thus, a persistent activation of *X* will trigger only a short burst of activity in *Z*, thus making this motif a “change detector” that returns to a default value after activation. Obviously, this intuitive description is critically dependent on the precise mathematical form of the interactions, and we describe two such forms next.

**Figure 1:**
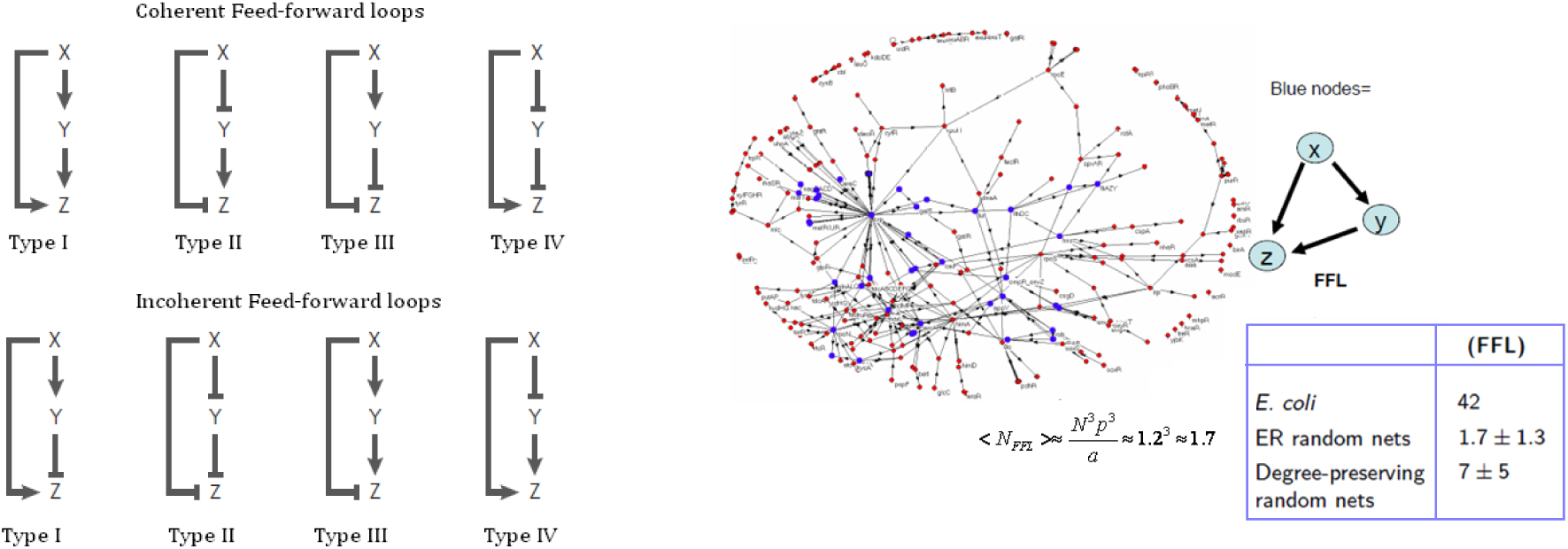
Left: four types of coherent (top) and incoherent (bottom) FFL’s. Blunt arrows ⊣ indicate repression, normal arrows → activation. Right: FFL’s are over-represented in *E. coli* transcriptional networks compared to random graphs. Both figures from [1].

We prefer to write the state of activation of the input mode *X* as *u* (as done in control theory for inputs), the activation of the output or reporter node *Z* as *Y*, and the intermediate or regulatory node *Y* as *x* (an internal state in control formalism). With these notations, the Type I IFFL can be written schematically as in the middle diagram in Fig. 2. The conceptual diagram may describe, in fact, various alternative molecular realizations. Different molecular realizations of the given motif can differ significantly in their dynamic response and, ultimately, biological function. Two realizations of the diagram in the middle of Fig. 2 are shown in that same figure. To be more precise, we study the simplest ordinary differential equation (ODE) models for these processes, in which the concentrations of the input *u* and species *x* and *y* are described by scalar time-dependent quantities. For the left reaction, we use the model

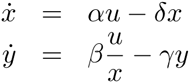

where dot indicates time derivative, the variables *u*, *x*, *y* and constants *α, β*, *δ*, γ are assumed positive, and the repression term 1/*x* is thought of as a quasi-steady state approximation of a binding event, or alternatively a Michaelis-Menten enzymatic reaction, that resulted in a term 1/(*K* + *x*) with *K* ≪ 1. For the right reaction, we use

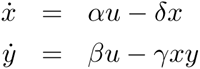

for suitable constants. For any given input function *u*(·) and initial values *x*(0) and *y*(0), the solution can be found by first solving the linear ODE for *x*(*t*), and then viewing the *y* equation as a linear ODE (with time-dependent coefficients). For constant inputs *u*(*t*) ≡ *u*_0_ *>* 0, both systems have the globally asymptotically stable steady state 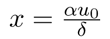 and 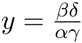. Observe that the steady state output value *y* is independent of the particular value of the constant input *u*_0_, meaning that both systems are *perfectly adapting* in the sense that the output approaches the same value irrespective of the input, provided that the input is constant. Of course, the transient behavior of *y* will depend on the initial conditions as well as the particular input being applied. For example, suppose that we consider a step signal that switches from *u*(*t*) = 1 to *u*(*t*) = 2 at time *t* = 5. Fig. 3(left) shows the brief activation behavior, followed by a return of the output to its adapted value. (In this and other simulations, we just take the constants *α* = *β* = *δ* = γ = 1, so that *y* = 1 at steady state.)

**Figure 2:**
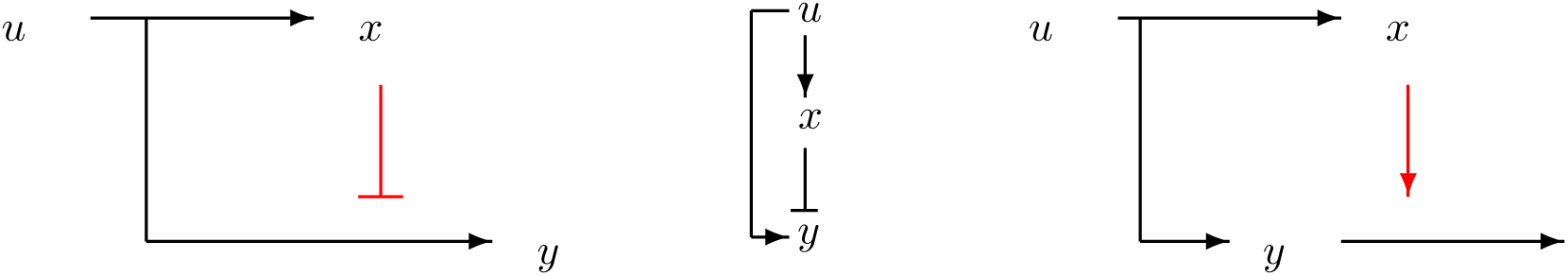
An incoherent feedforward loop (middle) and two instantiations, one through repression of production (left) and another one through enhancement of degradation (right). (Constitutive degradations of *x* and *y* not shown.)

**Figure 3:**
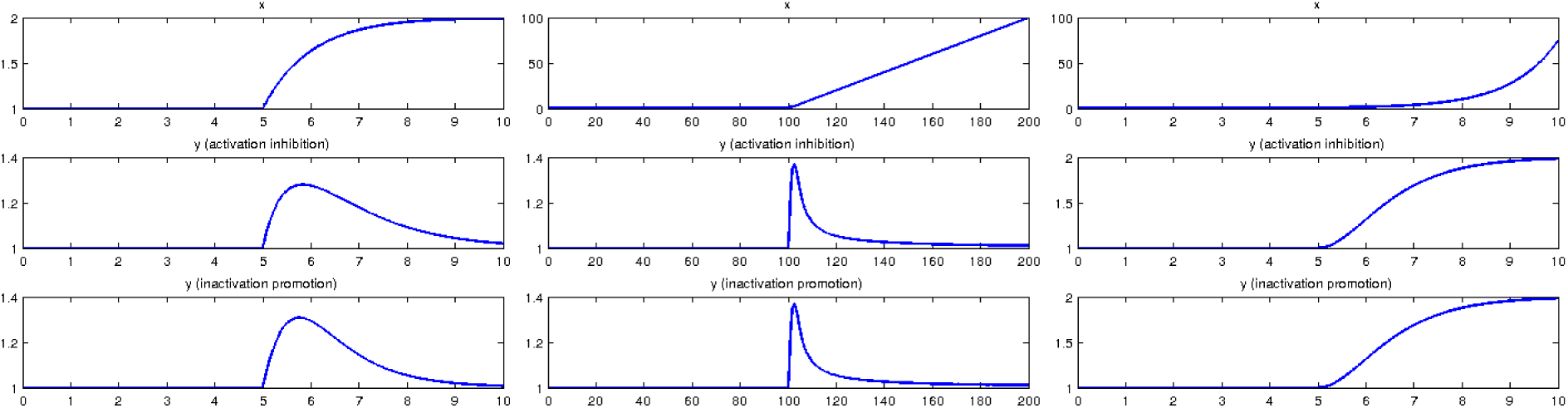
Simulations of both systems with *α* = *β* = *δ* = *γ* = 1. Initial conditions are *x*(0) = *y*(0) = 1. Horizontal axis is time (arbitrary units). Left: Step input *u* switches from *u*(*t*) = 1 to *u*(*t*) = 2 at time *t* = 5. After transient, output *y* returns to “hypoactivated” value 1. Middle: Input *u* switches from *u*(*t*) = 1 to *u*(*t*) = 1 +1, at time *t* = 100, Output converges again to “hypoactivated” value 1. Right: Input *u* switches from *u*(*t*) = 1 to *u*(*t*) = 1 + *t*, at time *t* = 5, with λ = 1. Value of output converges now to 1 + λ = 2.

In our context, we may think of the output *y* as the activation level of an immune system component, for example, a population if CD8+ effector T cells, systemically or in a particular microenvironment, and of *x* as the level of activation of a regulatory population, for example, a class of Treg cells. The input *u* is thought of as the level of presentation (by a specialized APC or perhaps a tumor) of a certain antigen.

For both of these model systems, the activation of the immune component will be at a “default” hypoactivated value, irrespective of the input u, provided that the activation signal *u* is constant. An increase in *u* will lead only to a brief spike of activity, unless the change persists.

Not only for constant inputs but also for “slowly varying” changes in activation there is only a spike in activity followed by return to default values: for linear growth, *y*(*t*) approaches its default value as *t* → ∞. An illustration is given in the middle panel of Fig. 3, which shows an input that switches from *u*(*t*) = 1 to *u*(*t*) = 1 + *t*. Convergence back to the default value of *y* is slower than for constant inputs, so we use a larger interval.

On the other hand, the response attains a larger “excitation” value when the input increases at a fast rate. For example, if the input switches from a constant level *u*(*t*) = 1 to an exponential growth *u*(*t*) = *e*^λ^*^t^*, *λ* > 0, then the system output *y*(*t*) approaches a value that exceeds its default (hypoactivated) value by an amount proportional to this rate of growth *λ*. Fig. 3(right) shows the switch to a more activated value. (For a larger rate *λ*, the effect will be more marked; we used a small value of *λ* to make the plots visually easier to compare.)

We also remark that periodic excitations leads to entrainment to excitations of the same frequency, see Fig. 4. This is consistent with Pradeu, Jaeger, and Vivier’s argument of how intermittent exposure to antigens might explain certain chronic autoimmune disorders.

**Figure 4:**
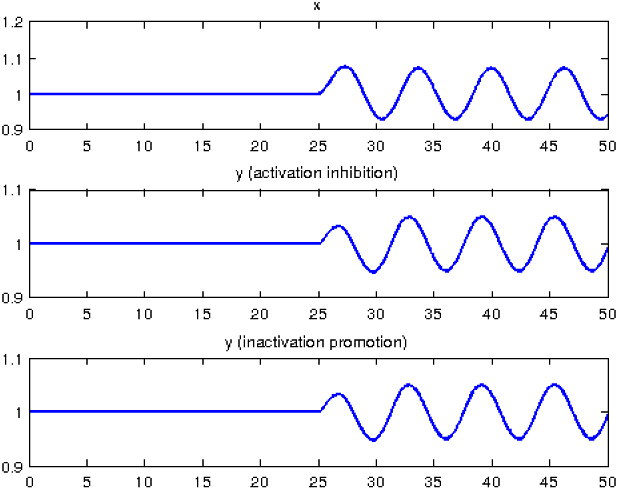
Simulations of both systems with *α* = *β* = *δ* = *γ* = 1, subject to periodic excitations. Input *u* switches from *u*(*t*) = 1 to *u*(*t*) = 1 + 0.1 sin *t* at time *t* = 25. Initial conditions are *x*(0) = *y*(0) = 1.

Obviously, our models so far have ignored the feedback of the immune system on the infection or tumor. Yet, even without the feedback effects, we have already some interesting and perhaps puzzling predictions. For example, IFFL change detectors would lead to a persistent immune excitation only for exponentially growing infection or tumor challenges. Any infection or tumor that achieves an equilibrium state (for example due to carrying-capacity limitations leading to logistic, Gompertz, etc., growth) would stop inducing an immune response.

A feature for future study is the addition of immune recruiting terms and positive feedback, perhaps by means of explicitly modeled cytokine and specifically chemokine signaling. Even a cascaded excitable system would lead to a permanent response if the signal from the IFFL detector is large enough in magnitude and duration.

### Treg cells as a possible regulatory node

Our analysis is merely a phenomenological “toy model” which does not specify immune components. Nonetheless, one might speculate that, as far as T cell activation and deactivation, regulatory T cells (Treg’s) may play a role as a regulating intermediate variable x. Treg’s are a type of CD4^+^ cell that play “an indispensable role in immune homeostasis” [6]. They express surface CD4 and CD25 and internally express the transcription factor FoxP3. Treg’s arise during maturation in the thymus from autoreactive cells (“natural Treg’s”), or are induced at the site of an immune response in an antigen-dependent manner (“induced Treg’s”). They are thought to play a role in limiting cytotoxic T-cell responses to pathogens, and Treg^−^ mice have been shown to suffer from extreme inflammatory reactions. It is known from animal studies that Treg’s inhibit the development of autoimmune diseases such as experimentally induced inflammatory bowel disease, experimental allergic encephalitis, and autoimmune diabetes [9]. Fig. 5 from [13] describes known mechanisms for Treg suppression of CTL’s.

**Figure 5:**
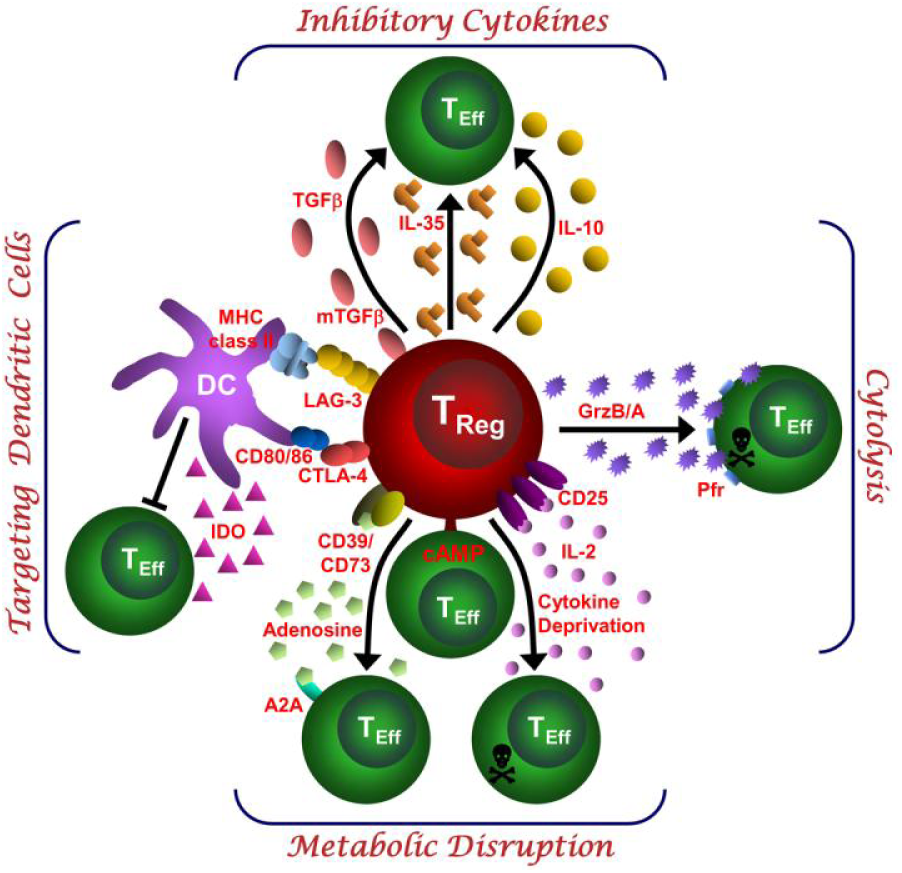
Schematic depicting the various regulatory T (Treg)-cell mechanisms arranged into four groups centered around four basic modes of action, from [13]. (1) Inhibitory cytokines. (2) Cytolysis. (3) Metabolic disruption. (4) Targeting dendritic cells.

## Perfect adaptation, Weber phenomenon, and scale-invariance

The property in which *y*(*t*) asymptotically returns to a pre-stimulus value under continuous stimulation is an example of the phenomenon called “perfect adaptation” in the theory of sensory systems, and is exhibited by systems processing light, chemical, and other signals. It has been extensively investigated both experimentally and mathematically [1, 7]. This notion can be refined in various ways, as we discuss next.

Suppose that we consider two step inputs *u*_1_ and *u*_2_ which are scaled versions of each other: *u*_2_(*t*) = *pu*_1_(*t*), for some positive number or scale factor *p*, as in Fig. 6(a). The perfect adaptation property requires that, whether excited by *u*_1_ or *u*_2_, the output signal will asymptotically return to the same value, as shown in Fig. 6(b). One stronger version of adaptation is the *Weber-Fechner* or “log sensing” requirement that, in addition, when starting from the corresponding adapted states, the maximum amplitude of the responses to the two scaled inputs should be the same, as illustrated in Fig. 6(c). A stronger property yet is the requirement that the two responses be identical as functions of time, again when starting from the corresponded adapted states. This last property is *scale invariance* or “fold change detection” of the response, and it was studied in some detail in [12, 11]. Recent interest in these properties was largely triggered by a pair of papers [4] and [3] published in late 2009, in which fold-change detection behavior was experimentally observed in a Wnt signaling pathway and an EGF pathway, respectively; these are highly conserved eukaryotic signaling pathways that play roles in embryonic patterning, stem cell homeostasis, cell division, and other central processes. Later, the paper [12] predicted scale invariant behavior in *E. coli* chemotaxis, a prediction which was subsequently experimentally verified [8].

**Figure 6:**
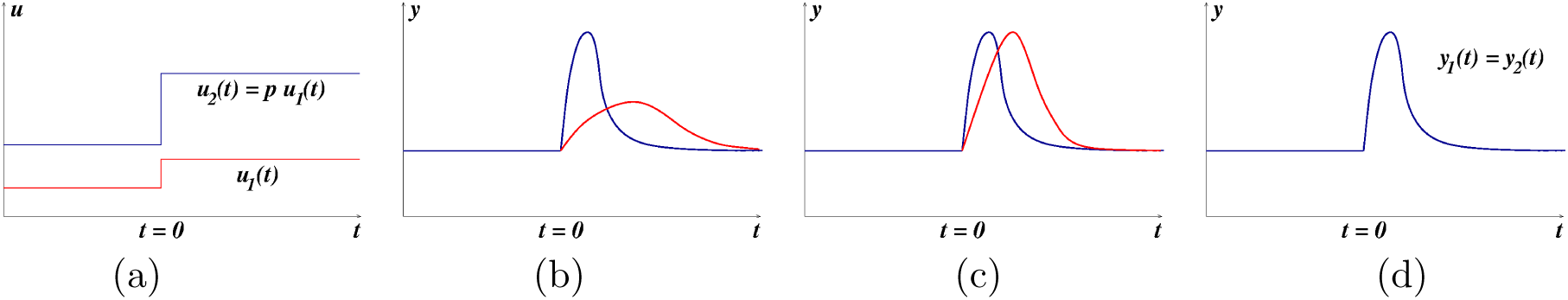
(a) Scaled step inputs and corresponding responses: (b) perfect adaptation; (c) Weber-like (same peak amplitude responses); (d) scale-invariance (same transient responses)

In our discussion of IFFLs, we introduced two circuits that implement Type I IFFLs (Fig. 2) and remarked that both lead to perfectly adapting systems. It is notable that only one of these exhibits scale invariance. Suppose that (*x*(*t*),*y*(*t*)) is any solution corresponding to the input *u*(*t*), for the system

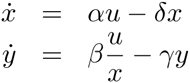

associated to Fig. 2(left). It is then immediate to verify that (*px*(*t*), *y*(*t*)) is a solution corresponding to the input *pu*(*t*), *t* ≥ 0, for any nonzero constant *p*. Thus, this system responds with the same output signal *y* (*t*) to two inputs that only differ in scale (provided that the initial state *x*(*t*) had already adapted to the input at time *t* < 0). In other words, given a step input that jumps from *u*(*t*) = *u*_0_ for *t* < 0 at time *t* = 0 and an initial state at time *t* = 0 that has been pre-adapted to the input *u*(*t*) for *t* < 0, *x*(0) = *αu*_0_ /*δ*, the solution is the same as when instead applying *pu*(*t*) for *t* > 0, but starting from the respective pre-adapted state *pαu*_0_ /*δ*. In contrast, scale invariance is false for the the degradation-induced inhibition described by the equations

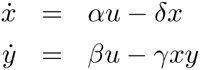

since multiplying *u* and *x* by *p* leads to a time-scale change in the output. See Fig. 7 for an example. Notwithstanding the extreme over-simplification represented by our model, it would be very interesting to test experimentally the response to scaled versions of antigen presentation, to help reverse engineer possible mechanisms.

**Figure 7:**
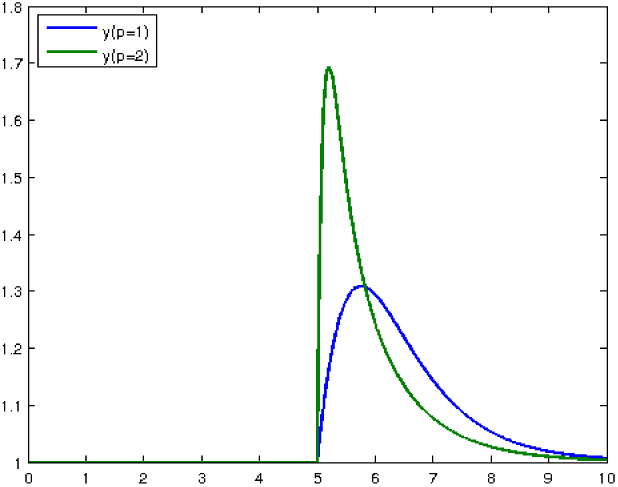
Simulations of degradation system with *α* = *β* = *δ* = *γ* = 1, comparing the responses to an input *u*(*t*) and an input *pu*(*t*), *p* = 10. Input *u* switches from *u*(*t*) = 1 to *u*(*t*) = 2 at time *t* = 5. Initial conditions are *x*(0) = *y*(0) = 1.

### A remark on feedback

Let us now add a term that affects the input (thought of as a pathogen load). We denote the pathogen population size (or a density in a particular environment) again by *u*(*t*), and assume that *u* evolves according to one of the following two possible differential equations:

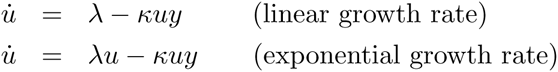

where λ and *κ* are positive constants. The mass-action term *κuy* is a killing effect due to the immune system. Reasoning informally, we may expect based on our previous discussion that in the linear growth case the immune reaction will be too weak to eliminate the pathogen: if *y* converges to its adapted value *y* = 1 (assuming for simplicity *α* = *β* = *δ* = *γ* = 1 as earlier) then we expect that *u*(*t*) ≈ λ – *κu* and *u*(*t*) might converge to λ/*κ*. On the other hand, in the exponential growth case, we expect based on our previous discussion that *y* will converge to some higher value, related to the effective growth rate of *u* (under feedback, so perhaps less than *λ*), and therefore *u* = (*λ* – *κy*)*u* means that, if *y*(*t*) > *λ*/*κ* for all *t*, then *u*(*t*)→ 0. Simulations agree with this informal reasoning, see Fig. 8. Thus, at least in this example, a pathogen that grows exponentially will be eliminated, but a slower linearly growing one might not. A follow-up paper will describe theory.

**Figure 8:**
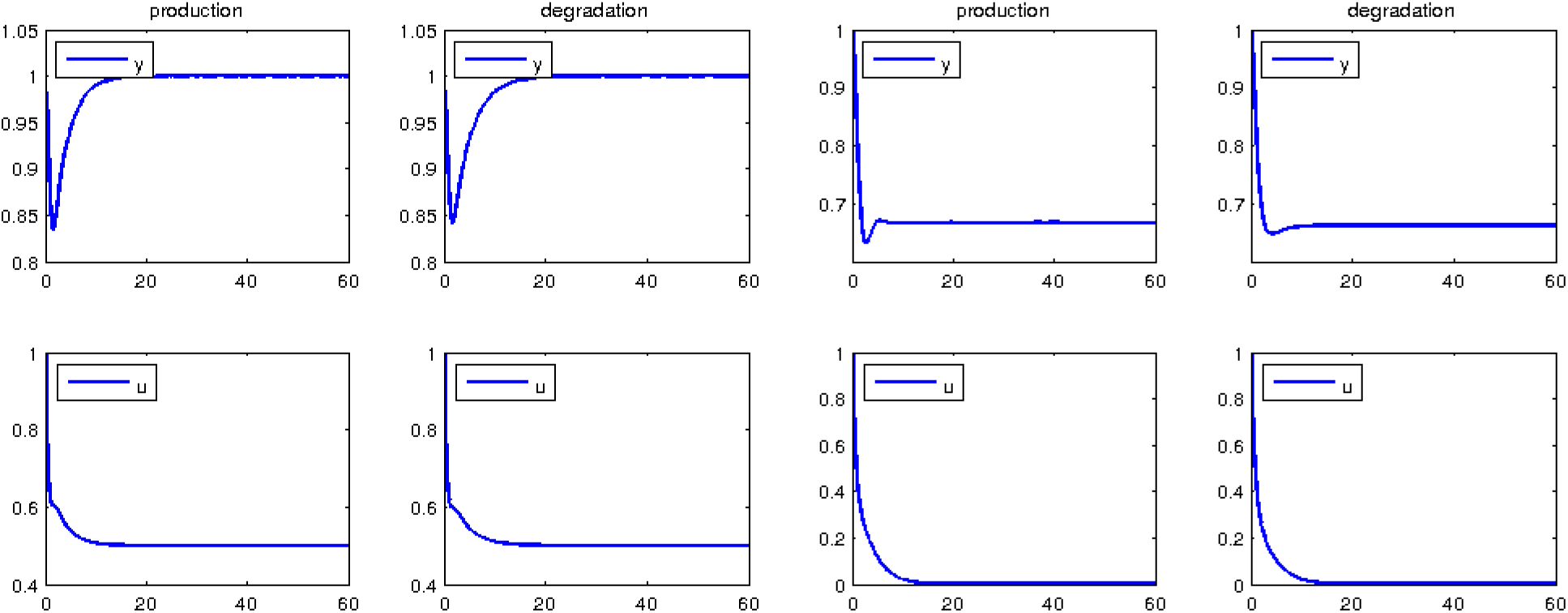
Simulations of the production inhibition and the degradation system, with *α* = *β* = *δ* = *γ* = 1 and λ = 1, *κ* = 2. Left two panels: linear growth. Right two panels: exponential growth.

## A few other models

We next briefly discuss our interpretation of some existing models for change detection in immune systems, and compare them to the IFFL model.

### Tunable Activation Threshold (TAT)

This model was suggested by Grossman and Paul [5] in 1992, motivated by the realization that “self/nonself discrimination may be much more complex than the simple failure of competent lymphocytes to recognize self-antigens”. The authors argued that for a stimulus to cause cell activation, the excitation level must exceed an activation threshold, and when engaged in persistent sub-threshold interactions, cells are protected against chance activation. In the TAT model, an *activation threshold* for an immune cell is dynamically modulated by an environment-dependent recent excitation history. This history is summarized by an *excitation index*, which we will denote as *x*(*t*), which computes a sort of weighted average of the cell’s past excitation levels. Given temporal excitation events, which we denote by *u*(*t*), it is assumed that the cell undergoes perturbations that depend on the difference between *u*(*t*) and the memory variable *x*(*t*). The key assumption is that such a perturbation, which we write as *y*(*t*):= *u*(*t*) − *x*(*t*), must exceed a fixed critical value, which we denote by *θ*, in order to cause activation. In other words, it must be the case that *u*(*t*) − *x*(*t*) > *θ*, or equivalently, *u*(*t*) > *x*(*t*) + *θ* (this is how we interpret the statement in [5] that “the activation threshold equals the excitation index plus that critical value”) for activation to occur. Cells maintained at a high level of excitation *x*(*t*) therefore are relatively insensitive to activation, thus being in some sense anergic. The authors deduce from their model that “upon gradual increase in the levels of excitation…a cell is not likely to be activated…it will become progressively anergic” which is intuitively equivalent to our remark about the lack of continued excitation under slow increases in antigen presentation. With our notations, the model suggested in [5] is:

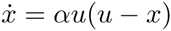

for some constant α, and the output would be *y* = *u* − *x*. (No explicit population-based nor signaling mechanism was given.) Notice that we then can derive a differential equation for y:

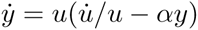

and this means, roughly, that *y* should approach 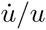, the logarithmic derivative of the input *u*, so that a “log sensing” property is satisfied by the output. Moreover, when the input is constant, the output converges to the same value (zero), independently of the actual value of the input, so we have perfect adaptation. Moreover, we expect *y* to be small (and thus not exceeding the threshold *θ*) unless *u* changes fast in the sense that its logarithmic derivative is large. For example, for *u*(*t*) increasing linearly, 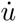 would be a constant, so 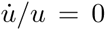 and therefore *y*(*t*) → 0 as *t* → ∞. On the other hand, for an exponentially increasing *u*(*t*), *y*(*t*) converges to a value proportional to the exponential rate. These properties are analogous to those satisfied by our model.

### Discontinuity theory of immunity

This model was suggested by Pradeu, Jaeger, and Vivier [10] in 2013 as a “unifying theory of immunity”. Their key hypothesis is that effector immune responses are induced by an “antigenic discontinuity” by which they mean a “sudden” modification of molecular motifs with which immune cells interact. The authors present evidence that natural killer (NK) cells and macrophages are activated by transient modifications, but adapt (ceasing to be responsive) to long-lasting modifications in their environment, and then propose to extend this principle to other components of the immune system, such as B cells and T cells. They also argue that although tumors give rise to effector immune responses, “a persistent tumour antigen diminishes the efficacy of the antitumor response”. In summary, their criterion of immunogenicity is the phenomenological antigenic discontinuity and not the nature of the antigen, including both “discontinuities” arising from self motifs such as tumors as well as from non-self motifs such as bacterial or viral infections. As examples of mechanisms for desensitization they mention receptor internalization, degradation or inactivation of signaling proteins. A concrete example of the latter is the dephosphorylation triggered by immunoreceptor tyrosine-based inhibition motif (ITIM)-containing receptors antagonizing kinases triggered by immunoreceptor tyrosine-based activation motif (ITAM)-containing receptors. The authors also mention Treg population dynamics.

Using our notations, the model in [10] starts by computing a running average of the absolute value of consecutive differences in inputs presented at discrete times on a sliding window *K* time units long:

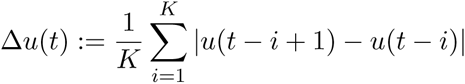

and then taking as the output *y* = Θ(Δ*u*), where Θ is a sigmoidal saturating function. The authors employ

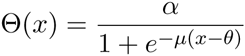

but one could equally well (and perhaps easier to justify mechanistically) employ a Hill-type function 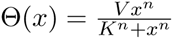. In continuous time, and assuming that the input is differentiable, we could interpret

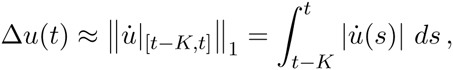

where the right hand term is the total variation of the input on this sliding window. Note the absolute value, which means that, in this model, activation is symmetrically dependent on increases or decreases of the excitation.^*^ As with our model, slow variations in the input will lead to small *y*(*t*), with the threshold function Θ resulting in an ultrasensitive, almost binary, response (provided that *μ* or *n* are large, in the two suggested functions Θ).

### Growth threshold conjecture

This model was suggested by Arias, Herrero, Cuesta, Acosta and Fernández-Arias [2] in 2015 as “a theoretical framework for understanding T-cell tolerance” based on the hypothesis that “T cells tolerate cells whose proliferation rates remain below a permitted threshold”. As in the other works, the authors postulate that T cells tolerate cognate antigens (irrespectively of their pathogenicity) as long as their rate of production is low enough, while those antigens that are associated with pathogenic toxins or structural proteins of either infectious agents or aggressive tumor cells are highly proliferative, and therefore will be targeted as foes by T cells. In summary, once again the postulate is that a strong immune response will be mounted against against fast-growing populations while slow-growing ones will be tolerated. The model in [2] is not one of change detection as such, but it is a closed-loop system that includes both detection and a killing effect on pathogens. To compare with our previous models, let us again denote the pathogen population size (or a density in a particular environment) by *u*(*t*) and the effector cell population by *y*(*t*). The authors give for *y* a second order equation 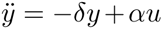, modeled on a spring-mass system that balances a “restoring to equilibrium force” to its activation by pathogens. We prefer to write the system as a set of first order ODE’s. Thus, we let 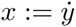, and write:

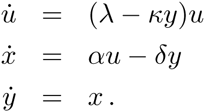

The *u* equation has an exponential growth term balanced by a kill rate that depends on the effector population. The effector population integrates the amount of *x* (which we might interpret as an intermediate type of cell); the growth of *x* is driven by pathogens, with a negative feedback from *y* (in essence an integral feedback on *x*), but there is no obvious biological mechanism for this model. Observe that when there is no pathogen, this results in a harmonic oscillator for *x* and *y*, with sustained oscillations and even negative values. In any event, the authors computationally obtain a bifurcation-like diagram in the (λ, *κ*) plane, dividing this plane into two regions, labeled “tolerance” (of infection, hence, failure of the immune system) and “intolerance”. These regions show how to trade off the growth rate λ of the pathogen versus the parameter *κ*, which represents a combination of affinity and clearance rate, and various conclusions regarding evasion strategies and the role of fever and even Treg cells are qualitatively derived from there.

## Footnotes

* Decreases may help with “missing self” recognition: the expression of a “self” marker suddenly decreases, triggering a response by NK cells and other immune components (Thomas Pradeu, personal communication).

